# Sulfide:quinone oxidoreductase drives mitochondrial supersulfide metabolism to regulate bioenergetics and longevity in eukaryotes

**DOI:** 10.64898/2026.04.05.716515

**Authors:** Jia Yao, Tetsuro Matsunaga, Akira Nishimura, Meg Shieh, Tomoaki Ida, Minkyung Jung, Seiryo Ogata, Tsuyoshi Takata, Uladzimir Barayeu, Hozumi Motohashi, Masanobu Morita, Takaaki Akaike

## Abstract

Sulfide:quinone oxidoreductase (SQR) is a critical enzyme that maintains sulfur metabolism by oxidizing sulfide to supersulfides, currently defined as sulfur metabolites with six valence electrons and no charge that are covalently catenated with other sulfur atoms and excludes disulfides. While SQR is known to contribute to mitochondrial electron transport, its physiological impact on systemic energy metabolism and longevity remains largely undefined. In this study, we investigated the role of SQR in mitochondrial bioenergetics and aging using SQR-deficient *Schizosaccharomyces pombe* (*Δhmt2*) and a mitochondria-selective SQR-deficient (*Sqrdl*^ΔN/ΔN^) mice model. Functional analysis demonstrated that *Δhmt2* grew normally in glucose but not in glycerol, indicating impaired mitochondrial respiration. It showed reduced membrane potential, ATP, and lifespan. Consistent with the yeast findings, *Sqrdl*^ΔN/ΔN^ mice exhibited accumulated levels of hydrogen sulfide and persulfides, and demonstrated impaired mitochondrial energy metabolism. Furthermore, supersulfide donor supplementation selectively conferred lifespan extension in wild-type yeast, but not in SQR-deficient strain, and similarly improved mitochondrial function exclusively in wild-type mouse embryonic fibroblasts, with no benefit observed in SQR-mutant counterparts. Together, our findings demonstrate that mitochondrial SQR plays an essential role in sulfur respiration, critically supporting mitochondrial function and organismal longevity across eukaryotes.

**Graphic Abstract:** 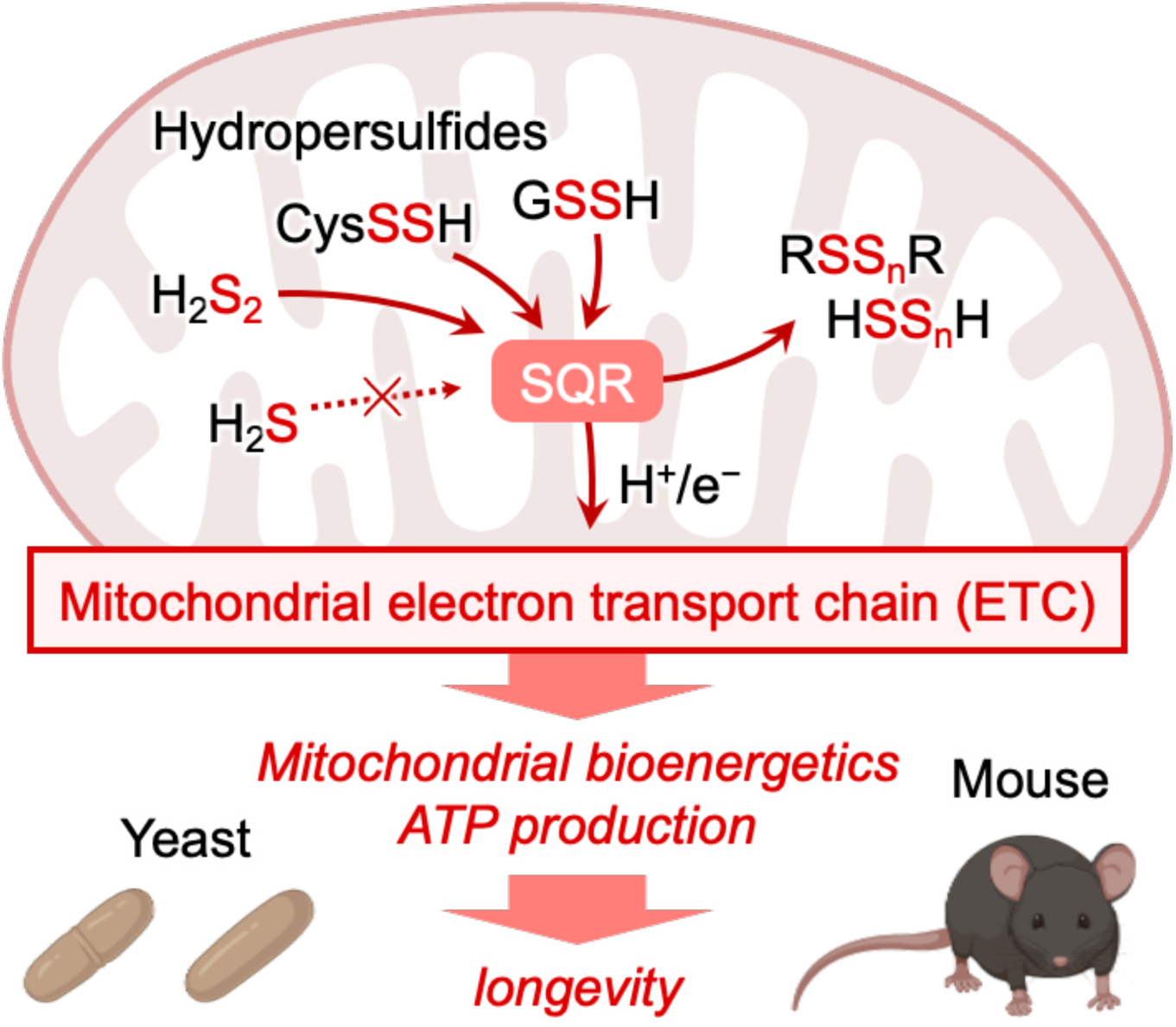

**Highlights:**

- Developed an SQR-deficient *S. pombe* (*Δhmt2*) model that exhibits sulfur metabolism, mitochondrial dysfunction, and shortened chronological lifespan
- Sulfide and supersulfide donors prolong yeast lifespan in a SQR-dependent manner
- Mitochondrial SQR is essential for membrane potential formation and ATP production in yeast and mammals

## 1. Introduction

Sulfur metabolism is a fundamental biochemical process that supports cellular signaling, redox homeostasis, detoxification, structural integrity, and energy production [1–3]. It involves the transformation of sulfur-containing compounds such as cystine, methionine, and hydrogen sulfide (H_2_S). Among these important sulfur species are supersulfides, which include hydropersulfides (RSSH) and polysulfides (RS_n_R, n > 2, R represents hydrogen or an alkyl group including cyclized forms). These species have recently emerged as key regulators of redox signaling and cellular functions [4–9]. Importantly, the dysregulation of supersulfides has been implicated in a wide range of diseases, including neurodegenerative and cardiovascular diseases, infectious diseases, skeletal abnormalities, and cancer, highlighting the importance of tightly regulated supersulfide metabolism [7, 10–15].

Sulfide:quinone oxidoreductase (SQR) is a key mitochondrial enzyme that catalyses the first committed step in sulfide oxidation by transferring electrons from H_2_S to ubiquinone in the electron transport chain (ETC) [16]. Originally identified in fission yeast as a determinant of cadmium (Cd) sensitivity [17], SQR is highly conserved across eukaryotes, including worms, fruit flies, mice, and humans [18, 19]. In mammalian systems, SQR has been primarily regarded as a detoxification enzyme that protects cells from sulfide toxicity [20–24]. However, most previous studies have relied on exogenous sulfide donors, leaving the physiological roles of SQR in endogenous sulfur metabolism mitochondrial function unknown. Emerging evidence suggests that SQR contributes to mitochondrial homeostasis by not only removing H_2_S, but also generating supersulfides intermediates that may participate in cysteine biosynthesis and redox regulation [25]. Nevertheless, the precise role of SQR in mitochondrial bioenergetics remains poorly understood.

The fission yeast *Schizosaccaromyces pombe* provides a unique model for studying SQR-dependent sulfur metabolism. Notably, *S. pombe* lacks key enzymes of the canonical cysteine biosynthesis pathway, including cystathionine β-synthase and cystathionine γ-lyase, as well as mitochondrial cysteinyl-tRNA synthetase (CARS2) and persulfide dioxygenase [4, 26]. Instead, *S. pombe* possesses a simplified mitochondrial sulfur metabolic pathway consisting primarily of SQR and 3-mercaptopyruvate sulfurtransferase. This minimal system enables direct investigation of SQR function in mitochondrial sulfur metabolism without confounding parallel pathways [27]. In this study, we generated an SQR-deficient *S. pombe* strain (*Δhmt2*) to investigate the role of SQR in mitochondrial function and organismal physiology. Using this model, we demonstrate that SQR is essential for mitochondrial energy metabolism, membrane potential formation, ATP production, and chronological lifespan in yeast.

We and others have shown that reactive persulfides such as cysteine hydropersulfide are primarily produced by cysteinyl-tRNA synthetase (CARS), a highly conserved enzyme now recognized for its moonlighting function as a cysteine persulfide synthase [6, 28]. In eukaryotes, the mitochondrial isoform CARS2 generates the majority of intracellular persulfides. We previously demonstrated that CARS2-dependent persulfide production is required to maintain mitochondrial membrane potential and ATP synthesis. Previous studies have reported that exogenously supplied sulfide increased oxygen consumption and mitochondrial membrane potential [25, 29], which implies that, under a particular condition, eukaryotes including mammals may exploit sulfide-based respiration as an evolutionary legacy derived from ancient organisms that lived in sulfide-rich habitats. For this paper, we hypothesized that SQR-mediated hydropersulfide oxidation is coupled with CARS2-mediated persulfide generation to fuel mitochondrial energy metabolism.

To investigate the physiological relevance of mitochondrial SQR-mediated sulfur metabolism in mammals, we employed a mitochondria-targeting-deficient SQR mutant mouse model. Through integrated sulfur metabolomics and bioenergetics in yeast, mouse embryonic fibroblasts (MEFs), and isolated mitochondria, we provide mechanistic evidence that SQR mediates electron transport and proton transfer from sulfide and hydropersulfide to the ETC. Collectively, our findings establish SQR as a central regulator of mitochondrial sulfur metabolism and reveal its critical role in driving sulfur-based energy metabolism and bioenergetics in eukaryotic cells.

## 2. Results

### 2.1. Validation of the Δhmt2 S. pombe model organism

Recent studies have indicated that the mitochondrial SQR is important for maintaining mitochondrial function by transferring electrons from H_2_S to coenzyme Q (CoQ), thereby linking sulfur metabolism to the ETC at the level of complex III [24, 25]. To investigate the role of SQR, we generated SQR-deficient fission yeast (*Δhmt2*) using the homologous recombination system (Fig. 1A). To verify the toxicity of CdCl_2_ towards the newly generated SQR-deficient yeast, various concentrations of CdCl_2_ (0 to 300 µM) were added to the culturing media of SQR-deficient and wild-type (WT) yeast and the subsequent growth rates were compared (Fig. 1B). As expected, the growth rate of the *Δhmt2* yeast significantly decreased with the increased concentrations of CdCl_2_. This was consistent with previously reported results [17], validating our model yeast for use in subsequent experiments. Additionally, the susceptibility of *Δhmt2* yeast towards NaHS (a sulfide donor) was also determined by comparing the growth rate of the *Δhmt2* yeast with that of the WT after being cultured in media containing 0-1,000 μM NaHS (Fig. 1C). As expected, *Δhmt2* growth rates decreased with increasing concentrations of NaHS compared to that of the WT, further validating our *in vivo* model.

**Fig. 1.**
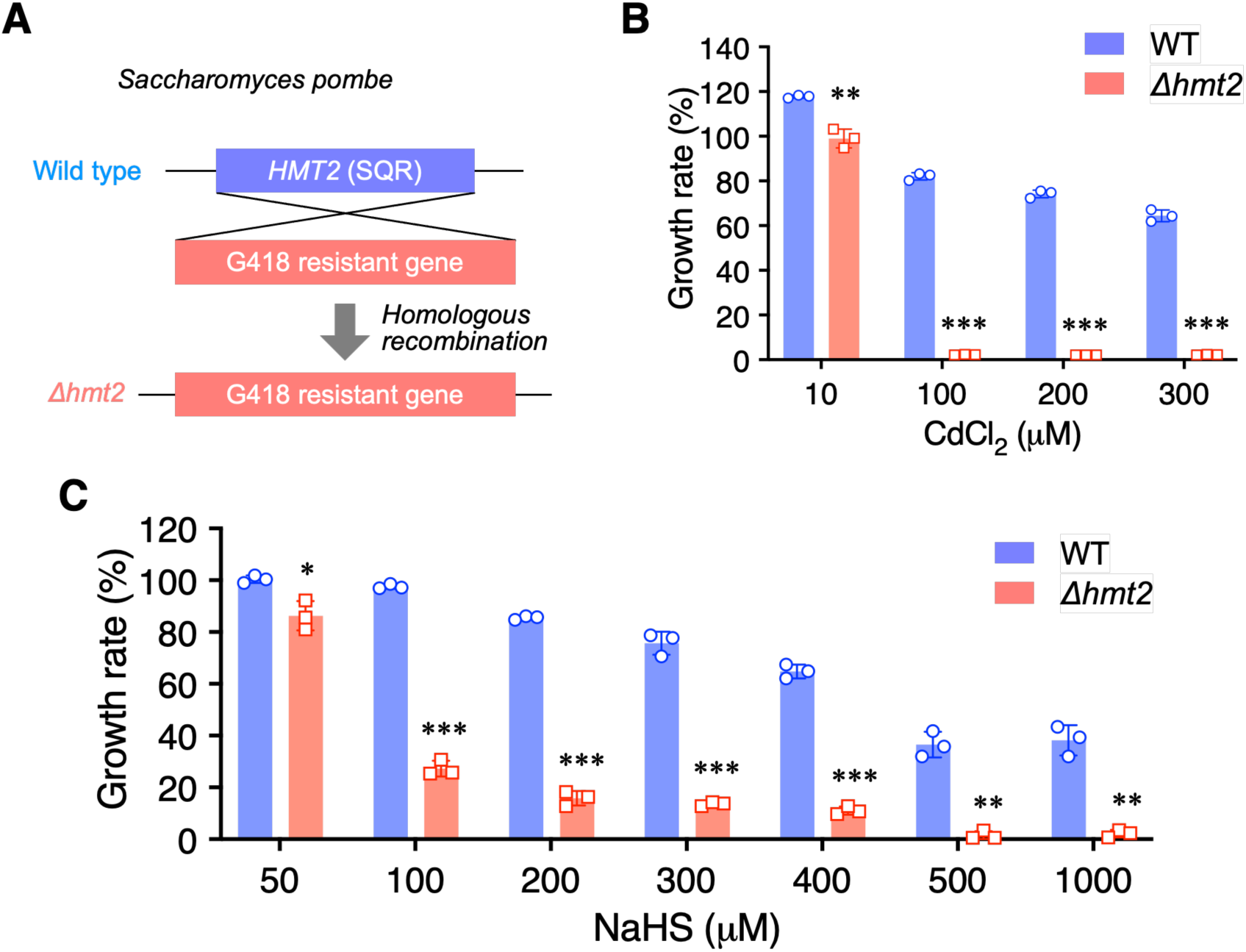
Construction of SQR mutant sensitive to hydrogen sulfide in fission yeast. (A) Construction of the *Δhmt2* strain. (B) Effect of CdCl_2_ treatment on yeast growth. Results are presented as means ± s.d. from three independent experiments, with statistical significance determined via Student’s *t*-test. \*\**P* < 0.01, \*\*\**P* < 0.001, *Δhmt2 vs.* WT. (C) Effect of NaHS treatment on yeast growth. Results are presented as means ± s.d. from three independent experiments, with statistical significance determined by Student’s *t*-test. \**P* < 0.05, \*\**P* < 0.01, \*\*\**P* < 0.001, *Δhmt2 vs.* WT.

### 2.2. Sulfur metabolism in SQR-deficient yeast

We next examined the impact of SQR deletion on sulfur metabolism and observed that SQR-deficient yeast (*Δhmt2*) had elevated hydrogen sulfide levels compared to that of WT yeast, as expected and consistent with the role of SQR in sulfide oxidation (Fig. 2). SQR deficiency also results in significantly reduced cysteine (CysSH) levels, consistent with a recently proposed role for SQR in cysteine biosynthesis (Fig. 2)[25]. However, SQR deficiency led to a greater accumulation of cysteine persulfide (CysSSH) and thiosulfate (HS_2_O_3_^-^) compared to that of WT yeast (Fig. 2), suggesting an important contribution of SQR in the oxidative metabolism of hydropersulfides in addition to its known role in sulfide oxidation.

**Figure 2.**
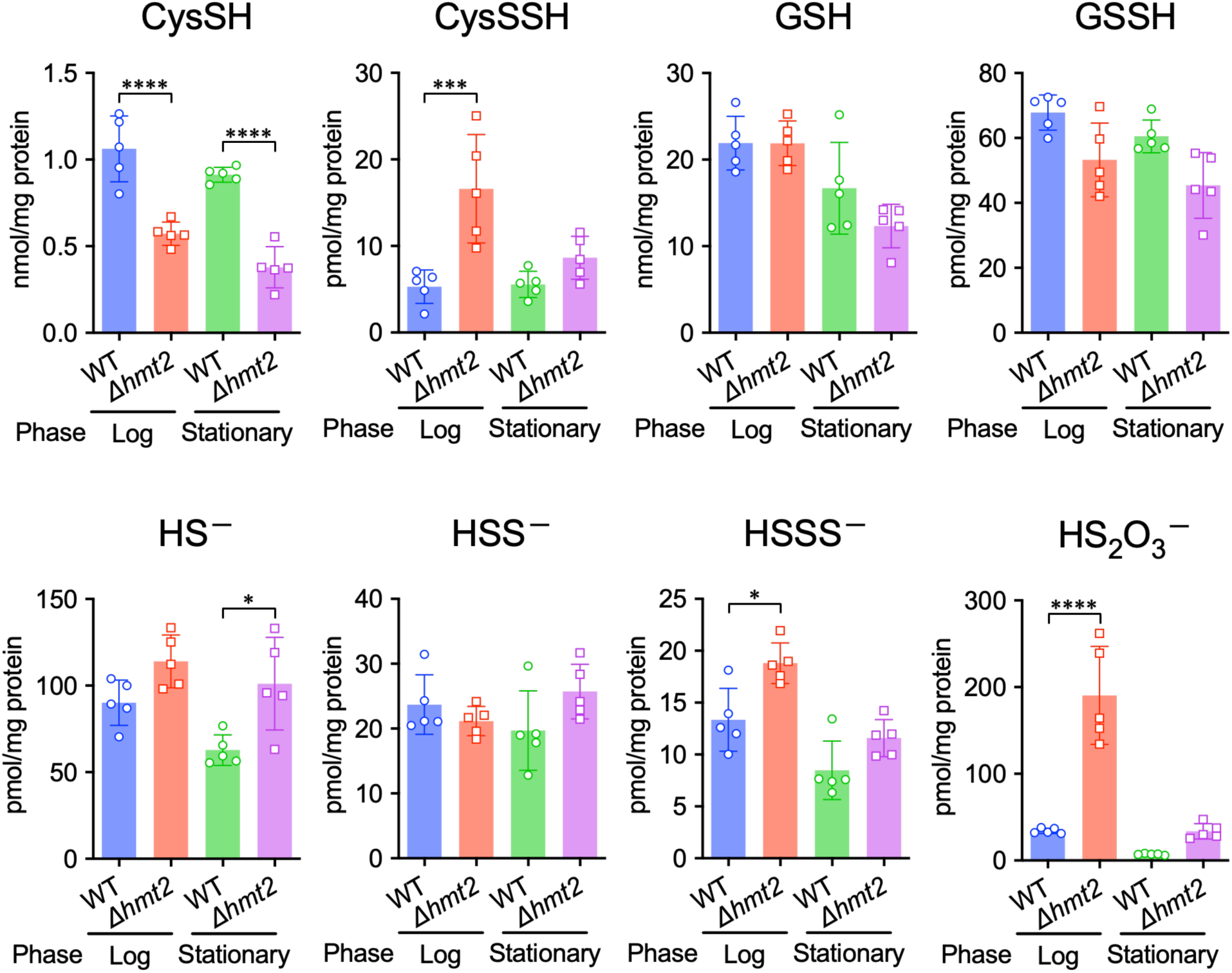
Mutation of SQR influences supersulfide metabolism in fission yeast. *In vivo* formation of various sulfur and supersulfide metabolites in the yeast WT and *Δhmt2* strain. Endogenous production of the analytes was identified through HPE-IAM labeling followed by LC-ESI-MS/MS analysis (n = 5). \**P* < 0.05, \*\*\**P* < 0.001, \*\*\*\**P* < 0.0001; Student’s *t*-test.

### 2.3. SQR is important in mitochondrial energy metabolism

To examine SQR’s contribution to mitochondrial energy metabolism, we compared yeast growth in either glycerol medium, where energy production during growth depends on mitochondrial energy metabolism, or in a glucose medium when energy production is primarily mediated by glycolysis and independent of mitochondrial respiration. Both WT and *Δhmt2* yeast grew similarly in glucose medium (Fig. 3A and B, left panels), but the growth of *Δhmt2* was strongly arrested compared with that of the WT yeast when glycerol was the sole carbon source (Fig. 3A and B, right panels). This finding suggests that SQR has an important role in mitochondrial energy metabolism. Moreover, we conducted JC-1 fluorescence staining with Fluorescence Activated Cell Sorter (FACS) analyses and observed increased mitochondrial membrane potential at the stationary phase when WT yeast were grown in glucose medium. This was not observed in SQR-deficient yeast (Fig. 3C, D), further demonstrating the importance of SQR in mitochondrial energy metabolism. Additionally, cellular ATP content was increased in WT yeast during stationary growth, while no such change was observed in that of SQR-deficient yeast (Fig. 3E). Thus, SQR-deficient yeast were shown to experience impaired mitochondrial energy production.

**Fig. 3.**
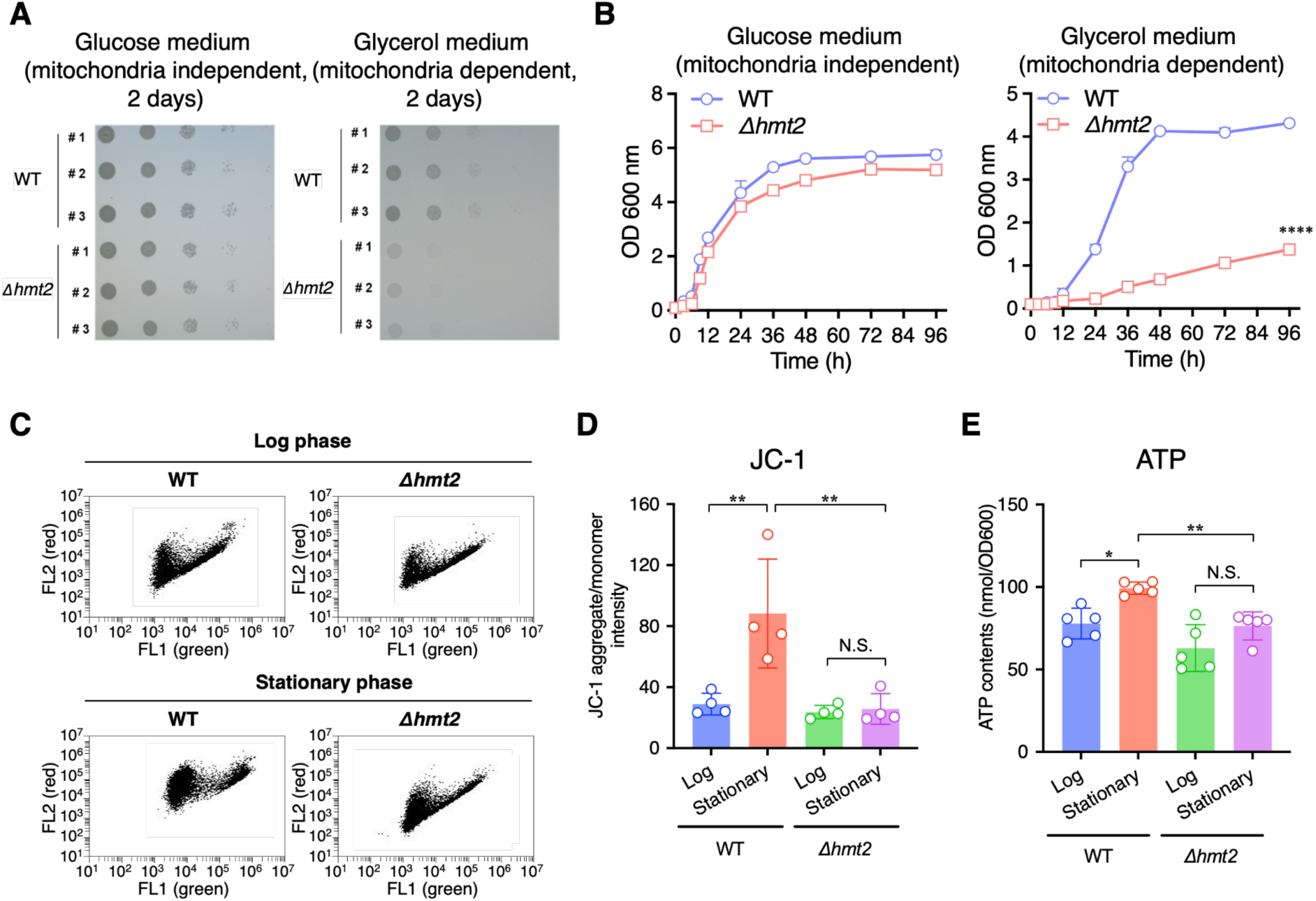
SQR promotes mitochondria-dependent energy metabolism in fission yeast. (A, B) Yeast growth test on solid medium (A) and liquid medium (B) in glucose medium (left panel) and glycerol medium (right panel), respectively. Data are presented as means ± s.d. from three independent experiments, with statistical significance determined by two-way ANOVA with Tukey’s test. \*\*\*\**P* < 0.0001, Δ*hmt2 vs.* WT. (C, D) Mitochondrial membrane potential evaluation by JC-1 at the log and stationary growth phases. Representative dot-plots of JC-1 flow cytometry (C), average fluorescence ratios of JC-1 aggregates (red) *vs.* JC-1 monomers (green) (D). Data are presented as means ± s.d. from four independent experiments. \*\**P* < 0.01, N.S., not significant, Δ*hmt2 vs.* WT, determined by one-way ANOVA with Tukey’s test. (E) ATP levels at the log and stationary growth phases. Data are presented as means ± s.d. from five independent experiments. \**P* < 0.05, \*\**P* < 0.01, N.S. (not significant), Δ*hmt2 vs.* WT, determined by one-way ANOVA with Tukey’s test.

### 2.4. SQR-deficiency decreases longevity

To determine the effect of the SQR deletion in our yeast model, we first precultured *Δhmt2* and WT yeast in SC medium for 3 days. Afterwards, cultures were diluted and plated on YPD plates to determine cell viability by measuring the efficiency of colony formation every three days (Fig. 4A). Our results demonstrated that WT yeast were able to form colonies for up to 18 days after culturing. On the other hand, *Δhmt2* yeast could only form colonies for less than 12 days, suggesting that the presence of SQR has a significant effect on the longevity of yeast.

**Fig. 4.**
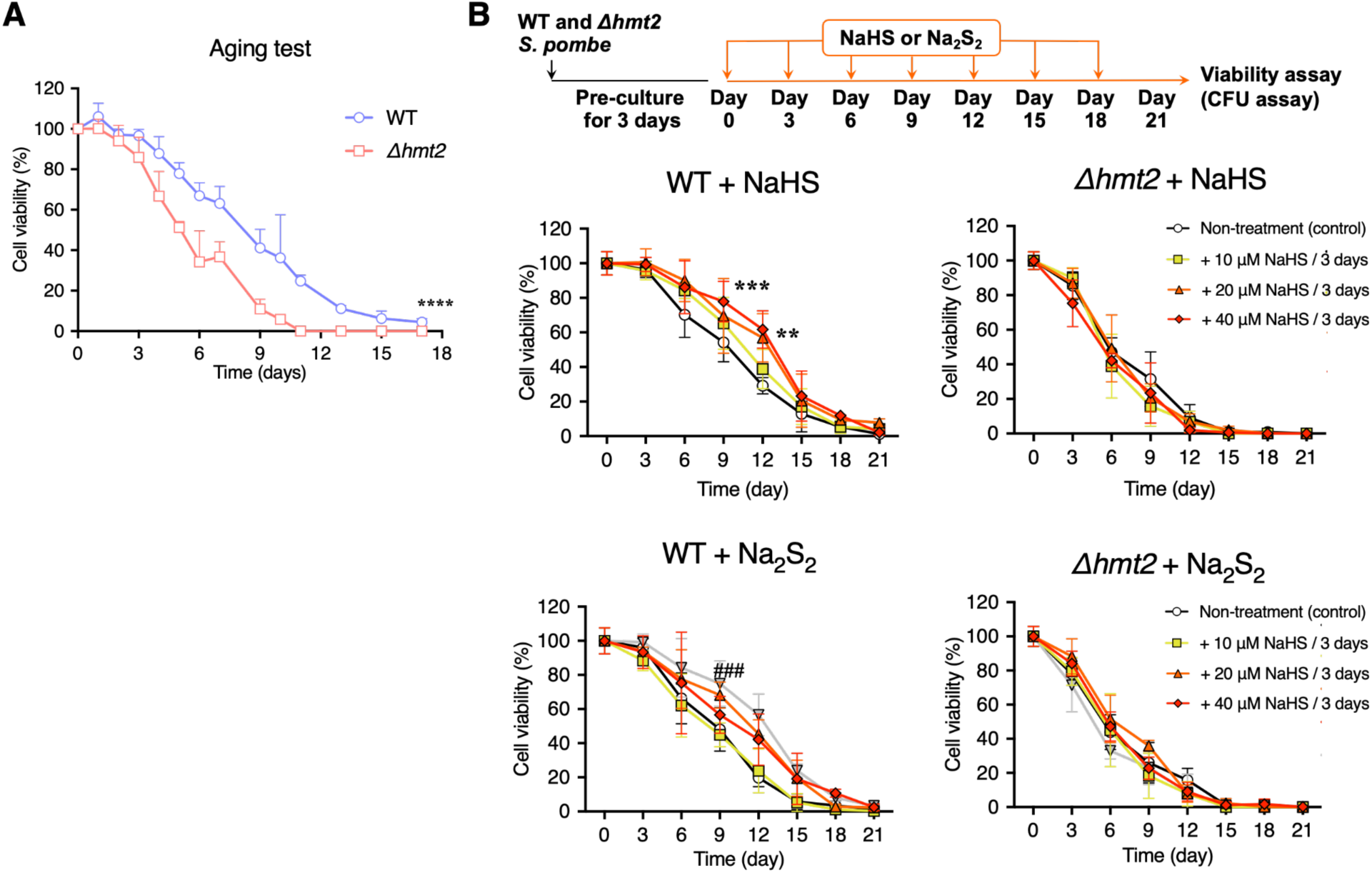
SQR contributes to chronological lifespan in fission yeast. (A) Chronological survival curves for WT and *Δhmt2* mutant yeast. Values are presented as means ± s.d. of five independent experiments. \*\*\*\**P* < 0.0001, Δ*hmt2* vs. WT, determined by two-way ANOVA with Tukey’s test. (B) Schedule of administration of sulfur donors. Time course of administration of NaHS or Na_2_S_2_ (upper panel). Chronological survival curves of yeasts supplemented with NaHS or Na_2_S_2_. Values are presented as means ± s.d. from five independent experiments. \*\**P* < 0.01 and \*\*\**P* < 0.001 for 40 µM NaHS treatment vs. control, respectively. ^###^*P* < 0.001 for 20 µM Na_2_S_2_ treatment *vs.* control, respectively. Significance was determined via two-way ANOVA with Tukey’s test.

### 2.5. Increased longevity with sulfur donors is SQR-dependent

Considering the role of SQR in oxidizing hydrogen sulfide to hydropersulfide and persulfide intermediates, we also wondered whether sulfide/persulfide donors could increase the lifespan of WT or SQR-deficient yeast (Fig. 4B). After treating the WT and *Δhmt2* yeast with the hydrogen sulfide donor NaHS or the hydropersulfide donor Na_2_S_2_ (the model equivalent of H_2_S_2_), we observed that both donors could extend the lifespan of WT yeast by at least 3 days. On the other hand, the donors did not affect the lifespan of SQR-deficient yeast (Fig. 4B). This result suggests that SQR can utilize both hydrogen sulfide and hydropersulfides as substrates to promote the lifespan extension of *S. pombe*, which could potentially thereby influence the bioenergetics and maintenance of the mitochondrial membrane potential in the yeast’s mitochondria. As *Δhmt2* yeast were SQR-deficient, their ability to utilize sulfides and persulfides was impacted, likely resulting in the lack of effect of the donors on extending the lifespan of the yeast.

### 2.6. Mitochondrial SQR deficiency in mice causes emaciation and premature death after weaning

To investigate the physiological relevance of mitochondrial SQR-mediated sulfur metabolism, we developed mitochondrial SQR-deficient mice that expressed SQRΔN, an N-terminally truncated SQR that lacks the mitochondrial targeting sequence (Fig. S1)[29]. By using CRISPR-Cas9 technology, we disrupted the ATG start codon of the murine *Sqrdl* gene by a 14-bp deletion, resulting in an in-frame translation from a downstream ATG and production of an N-terminally truncated SQR lacking its mitochondrial localization signal; consequently, the SQR protein failed to localize to the mitochondria (Fig. S1A)[29].

### 2.7. Mitochondrial SQR deficiency results in persulfide accumulation

In yeast, SQR-mediated effects on mitochondrial metabolism and longevity were closely correlated with the role of SQR in the metabolism of H_2_S and CysSSH. Therefore, we decided to study the overall effects of mitochondrial SQR deficiency or mislocalization of SQR to the cytoplasm on sulfur metabolism and levels of supersulfides in *Sqrdl*^ΔN/ΔN^ immortalized (MEFs)s and their isolated mitochondria (Fig. 5 and Fig. S2A). In the *Sqrdl*^ΔN/ΔN^ iMEFs, we found a dramatic increase in levels of hydropersulfides, including cysteine hydropersulfide, glutathione hydropersulfide, and hydrogen persulfide (HSSH) and H_2_S. This suggests that not only H_2_S but also hydropersulfides, such as cysteine hydropersulfide and glutathione hydropersulfide, serve as primary substrates for SQR. This is because SQR inactivation can result in persulfide accumulation; the same phenotype was observed in our yeast models.

**Fig. 5.**
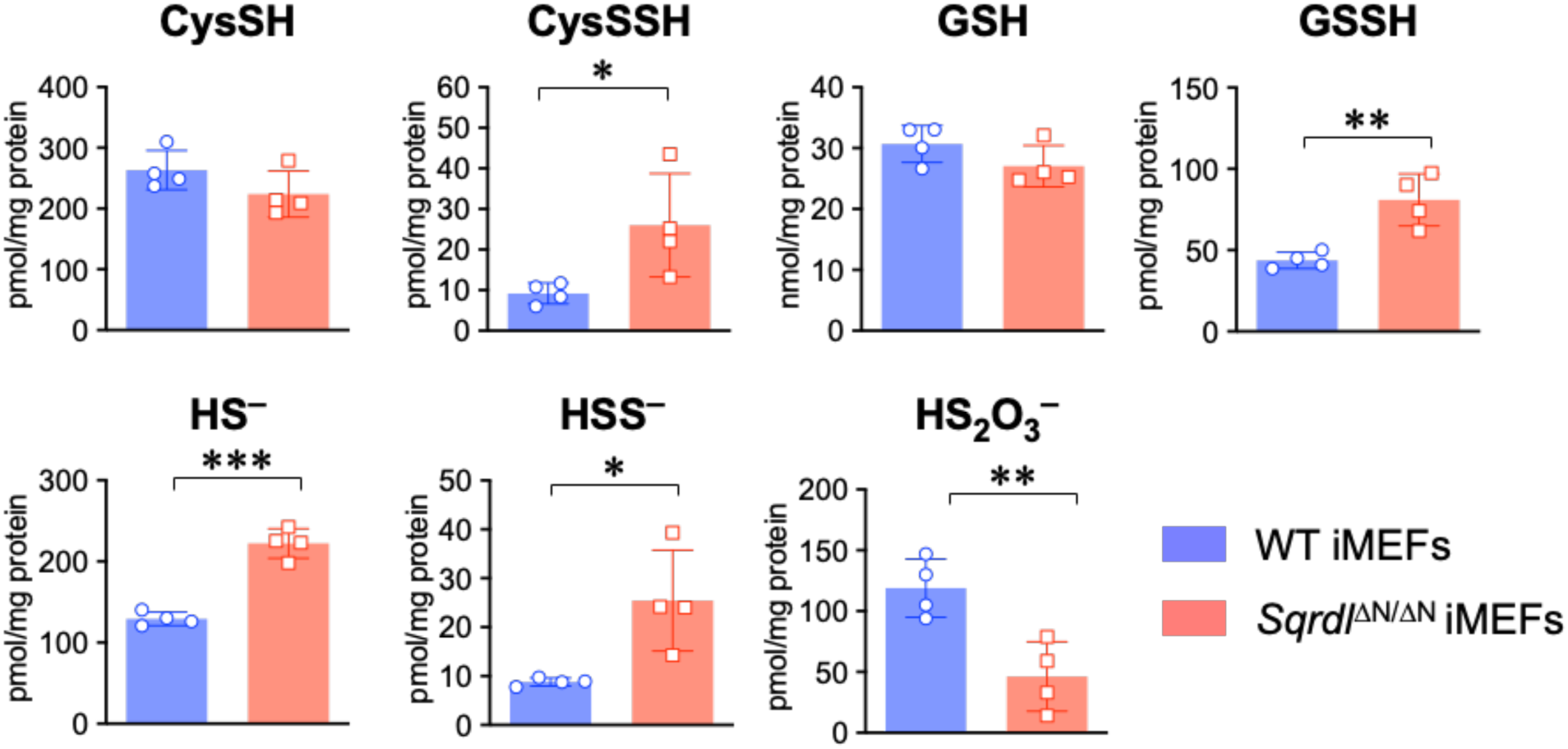
Sulfur metabolome analysis for SQR mutant mice. Intracellular levels of sulfur metabolites in WT and *Sqrdl*^ΔN/ΔN^ iMEFs. Data are means ± s.d. (*n* = 4). \**P* < 0.05, \*\**P* < 0.01 and \*\*\**P* < 0.001, determined by Student’s *t*-test.

### 2.8. Mitochondrial SQR deficiency attenuates mitochondrial membrane potential

Our data in *S. pombe* and iMEFs strongly suggests that SQR is required for the mitochondrial bioenergetics of eukaryotes under physiological conditions, and that inactivation of mitochondrial SQR results in the accumulation of H_2_S and supersulfides. On the basis of our previous studies [6, 30] and other studies [31, 32], we hypothesized that SQR oxidizes the hydrogen sulfide and hydropersulfide that are generated in the mitochondria and contributes to mitochondrial membrane polarization by transferring electrons and protons to ubiquinone in the ETC. To investigate this hypothesis in eukaryotes, we studied the effects of mitochondrial SQR elimination on the cellular energy metabolism of iMEFs due to the difficulties associated with isolating mitochondria from yeast. Specifically, iMEFs were derived from *Sqrdl*^ΔN/ΔN^ mice, in which the SQR protein is expressed but fails to translocate into the mitochondria. JC-1 fluorescence imaging was used to assess the mitochondrial membrane potential formation (Fig. 6A). The mitochondrial membrane potential was found to be reduced in *Sqrdl*^ΔN/ΔN^ iMEFs in comparison to that of WT iMEFs under standard culturing conditions (Fig. 6B). On the other hand, the oxygen consumption rate (OCR) of *Sqrdl*^ΔN/ΔN^ iMEFs was similar to that of the WT iMEFs (Fig. 6C), indicating that electron transport is not significantly affected in *Sqrdl*^ΔN/ΔN^ iMEFs. These results suggest that proton translocation per oxygen consumption is decreased in the absence of mitochondrial SQR. Meanwhile, increasing the concentrations of cystine, NaHS, and Na_2_S_2_ in the culturing media of WT iMEFs all showed SQR-dependent elevation of the membrane potential (Fig. 6D-F). These results suggest that proton/electron donation from hydrogen sulfide and hydrogen persulfide to ubiquinone as mediated by SQR efficiently generates the membrane potential. We also closely analyzed the contribution of SQR to the generation of the mitochondrial membrane potential by utilizing specific inhibitors of ETC components. To amplify the membrane potential, we used carboxyatractyloside (CATR), an inhibitor of mitochondrial ADP/ATP translocase [33]. CATR treatment increased the membrane potential in WT iMEFs, but this increase in *Sqrdl*^ΔN/ΔN^ iMEFs (Fig. S3A) was not observed, suggesting that the proton supply for the proton gradient was reduced in the *Sqrdl*^ΔN/ΔN^ iMEFs. The CATR-induced increase in membrane potential in the WT iMEFs was also not significantly affected by rotenone and atpenin A5, inhibitors of complex I and complex II, respectively, but was reduced by antimycin A, an inhibitor of complex III (Fig. S3B). Taken together, these results suggest that SQR is involved in the proton supply for maintaining mitochondrial membrane potentials independent of complexes I and II but is dependent on complex III. Thus, SQR may mediate proton donation from hydrogen sulfide and hydrogen persulfide to ubiquinone, which is located downstream of complexes I and II and upstream of complex III (Fig. S3C).

**Fig. 6.**
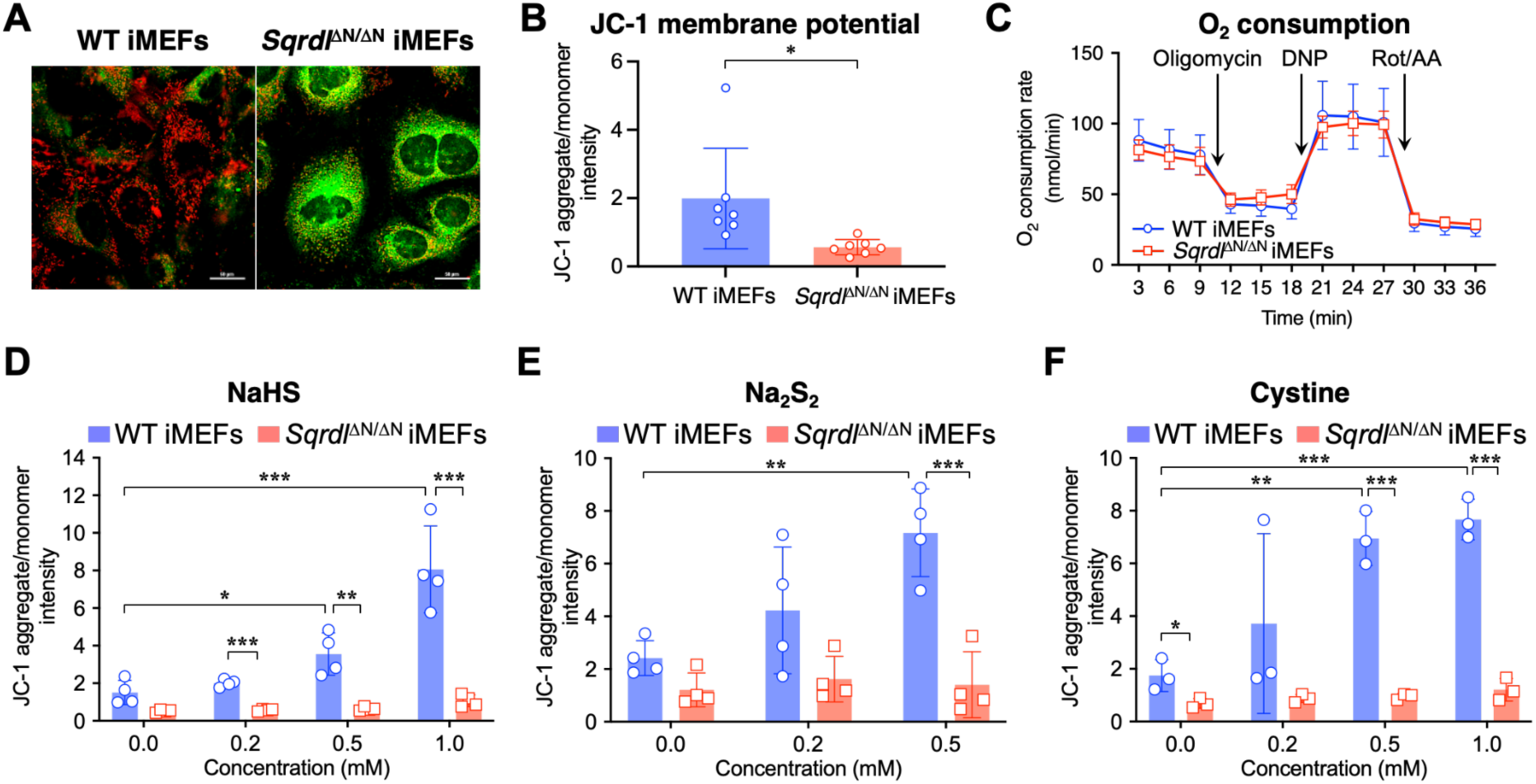
Whole-cell evaluation of SQR’s contribution to mitochondrial activity. (A) Schematic illustration(A) and measurement (B) of JC-1 iMEFs staining, with representative confocal fluorescence images of JC-1-stained WT and *Sqrdl*^ΔN/ΔN^ iMEFs. Scale bars, 50 µm. (C) Measurement of the OCR in WT and *Sqrdl*^ΔN/ΔN^ iMEFs. The OCR was measured with the XF96 Extracellular Flux Analyzer. Data are represented as mean ± s.d. (*n* = 10). Rot, rotenone; AA, antimycin A; DNP, 2,4-dinitrophenol. (D-F) Effects of the addition of NaHS (D), Na_2_S_2_ (E), and cystine (F) on mitochondrial membrane potential in WT and *Sqrdl*^ΔN/ΔN^ iMEFs. Data shown in (D-F) are mean ± s.d. (*n* = 3 or 4). \**P* < 0.05, \*\**P* < 0.01, \*\*\**P* < 0.001, N.S. (not significant), determined by one-way ANOVA with Tukey’s test.

### 2.9. Hydrogen sulfide and hydrogen persulfide generate mitochondrial membrane potential in an SQR-dependent manner

To more closely analyze the direct effects of hydrogen sulfide and hydropersulfide on mitochondrial membrane potential and complex I-IV activity, we established a tetramethylrhodamine ethyl ester (TMRE) time-lapse imaging system of isolated mitochondria from iMEFs (Fig. S4A-C). Sulfide supplementation to isolated mitochondria was found to generate the membrane potential in an SQR-dependent manner, although the effective sulfide dose was much lower for the isolated mitochondria than for the whole cells (Fig. 7A and Fig. S4D). The addition of 3 μM NaHS retained the membrane potential response to succinate, malate, and glutamate and to *N,N,N’,N’*-tetramethyl-1,4-benzenediamine (TMPD) with ascorbate, which indicates that sulfide does not suppress the activities of complex I/II and complex IV. In contrast, the membrane potential response was halted by concentrations of NaHS above 30 μM, suggesting that complex IV is sensitive to sulfide-mediated inhibition (data not shown). The sulfide-mediated generation of membrane potential was not affected by the inhibition of complexes I and II (Fig. 7B) but was stopped by complex III inhibition (Fig. 7C).

**Fig. 7.**
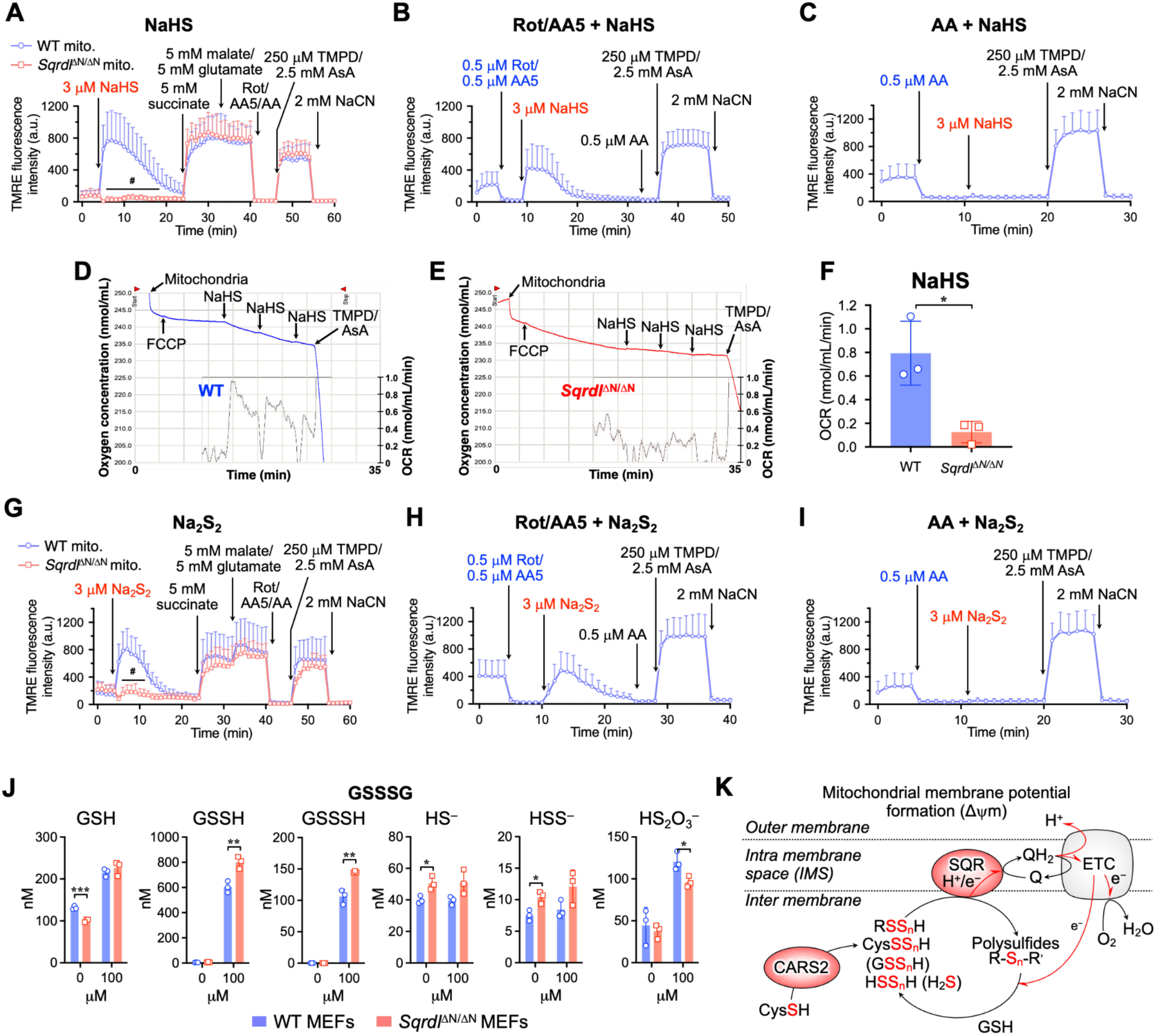
Evaluation of the SQR contribution to mitochondrial activity in isolated mitochondria. (A-C) The membrane potential of isolated mitochondria was monitored by means of the TMRE fluorescence intensity. Mitochondria isolated from WT and *Sqrdl*^ΔN/ΔN^ iMEFs were treated with NaHS (A) followed by treatments with succinate/glutamate/malate, Rot/AA5/AA, TMPD/AsA, and NaCN. Data are shown as means ± s.d. AsA, ascorbic acid. ^#^*P* < 0.0001, determined by two-way ANOVA with Tukey’s test. (B, C) NaHS-induced membrane potential of mitochondria isolated from WT iMEFs in the presence of 0.5 µM Rot/0.5 µM AA5 (B) and 0.5 µM AA (C). Data are shown as mean ± s.d. Representative oxygen consumption in response to sulfide by mitochondria isolated from mouse livers of WT mice (D) and *Sqrdl*^ΔN/ΔN^ mice (E); data are from three independent experiments. (E) Oxygen consumption in response to NaHS by mitochondria isolated from the livers of WT and *Sqrdl*^ΔN/ΔN^ mice. Data are means ± s.d. of average OCR induced by NaHS in three independent experiments. \**P* < 0.05 as determined by Student’s t-test. (G-I) Mitochondria isolated from WT and *Sqrdl*^ΔN/ΔN^ iMEFs were treated with Na_2_S_2_ (G) followed by treatments with succinate/glutamate/malate, Rot/AA5/AA, TMPD/AsA, and NaCN. Data are shown as mean ± s.d. ^#^*P* < 0.0001 as determined by two-way ANOVA with Tukey’s test. (H,I) Na_2_S_2_-induced membrane potential of mitochondria isolated from WT iMEFs in the presence of 0.5 µM Rot/0.5 µM AA5 (H) and 0.5 µM AA (I). Data are shown as mean ± s.d. (J) Sulfur metabolites in mitochondria isolated from WT and *Sqrdl*^ΔN/ΔN^ iMEFs with or without GSSSG supplementation. Data are represented as mean ± s.d. (*n* = 3). \**P* < 0.05, \*\**P* < 0.01, \*\*\**P* < 0.001, determined by Student’s *t*-test. (K) A schematic model of SQR-driven sulfur respiration in the mitochondria. Hydropolysulfides (shown as RSS_n_H) synthesized by CARS2 are oxidized by SQR and transformed into polysulfides (R-S_n_-Rʹ, *n* ≥ 2), which are coupled with the generation of membrane potential. The resulting R-S_n_-Rʹ is expected to accept electrons and protons from the ETC and the thioredoxin/thioredoxin reductase system (Trx/TrxR) in the mitochondria. Q, ubiquinone; QH_2_, ubiquinol; NADPH, nicotinamide adenine dinucleotide phosphate reduced; ΔΨm, membrane potential.

Considering these results, we wondered whether they could also be observed in diseased human mitochondria. The translatability of our results from iMEFs was promising, so we next sought to verify the electron transport from our sulfide donor in a more complex model. Mitochondria were isolated from the livers of WT and *Sqrdl*^ΔN/ΔN^ mice, and their oxygen consumption in response to sulfide treatment was evaluated. A 10 μM NaHS supply induced a small oxygen consumption only in WT mitochondria (Fig. 7D-F), suggesting that the SQR was responsible for the sulfide-derived electron transport. The oxygen consumption induced by the NADH-generating substrates malate and glutamate, followed by a supply of ADP, showed no significant differences between WT and *Sqrdl*^ΔN/ΔN^ mitochondria (Fig. S4E, F), results that were consistent with the whole-cell OCR measured in normal culture media (see Fig. 6C). Because hydropersulfides may also be substrates for SQR (see Figs. 5–7), we also investigated the effect of hydropersulfides on mitochondrial membrane potential. The addition of Na_2_S_2_ generated results that were quite similar to that of those induced by NaHS (Fig. 7G-I), suggesting that hydropersulfides can serve as comparably favorable substrates for SQR. It must be noted, however, that despite being widely used in the field as a model equivalent of H_2_S_2_, Na_2_S_2_ is a fast, uncontrollable donor that releases H_2_S_2_ in conjunction with H_2_S_n_ and H_2_S. To further dissect the contribution of different sulfur species to mitochondrial membrane potential formation, we performed experiments in the presence of an H₂S scavenger (SS19)[34]. Notably, even under SS19 pretreatment conditions, NaHS efficiently induced mitochondrial membrane potential formation (Fig. S4G). Likewise, hydrogen persulfide (Na₂S₂) robustly generated membrane potential in the presence of SS19 (Fig. S4H). These findings suggest that the bioenergetic effects observed are not mediated by free H₂S itself but rather by downstream supersulfide, most likely hydropersulfides and polysulfides. Indeed, the addition of oxidized glutathione trisulfide (GSSSG) to isolated mitochondria increased levels of glutathione hydropersulfide (GSSH), which in turn is likely to serve as a substrate for SQR (Fig. 7J). Taken together, our data suggest a model in which H_2_S and supersulfides can act as substrates for SQR that feeds electrons into the ETC to drive oxidative phosphorylation under conditions of elevated sulfur metabolism (Fig. 7K).

## 3. Discussion

This study demonstrated the essential functions of SQR in promoting mitochondrial persulfide synthesis and in maintaining metabolic activities. Our results suggest that eukaryotes (*i.e.* yeast, mammals, etc…) utilize sulfur respiration under physiological conditions. Interestingly, both sulfide and hydropersulfides could be oxidized by SQR, leading to electron and proton donation to the ETC via ubiquinone and maintaining the mitochondrial membrane potential. This finding is a missing piece of the sulfur respiration system that we proposed in our previous study [6, 30]. Hydropersulfides produced by CARS likely serve as substrates for SQR, as they can donate electrons and protons to CoQ, and the resulting oxidized persulfides are expected to receive electrons from the ETC and other redox systems in the mitochondria. From this, a new aspect of energy metabolism through sulfur respiration has now updated the concept of cysteinolysis [35]; cysteine is a source of hydropersulfides, which play a central role not only in cellular redox reactions, but also in mitochondrial energy metabolism because they can serve as substrates for ATP production.

In our previous study, we proposed that cysteine hydropersulfide synthesized by CARS2 served as an electron acceptor from the ETC and that the resulting hydrosulfide served as a substrate for SQR, thus generating the mitochondrial membrane potential [6, 29]. Consistent with this model, CARS-deficient yeast exhibited decreased production of cysteine hydropersulfide and other supersulfides, accompanied by an impaired mitochondrial energy metabolism [36]. In our current study with *S. pombe*, however, SQR deficiency in the mitochondria unexpectedly resulted in a large accumulation of hydropersulfides as well as hydrogen sulfide, which strongly suggests that both hydropersulfides and hydrogen sulfide are direct substrates for SQR (Fig. 2). Similar results were reported from the analysis of SQR mutant mice (Figs. 5–7 and S2, 4), further supporting the growing role of SQR in the metabolism of both hydropersulfides and hydrogen sulfide [29]. This result allowed us to revise our model of sulfur respiration, where cysteine hydropersulfide and derivative hydropersulfides (e.g. GSSH) generated by CARS2 are oxidized by SQR and serve as electron and proton donors in the ETC. The resulting oxidized persulfides (most typically, GSSSG) function as electron acceptors in the ETC. Thus, the idea that sulfur respiration in eukaryotes can operate under physiological conditions to exploit the redox cycles of persulfides is highly plausible. Additional detailed studies are required to obtain solid evidence that persulfides can act as electron acceptors equivalent to oxygen.

Interestingly, our scavenger experiments further refine this model. The persistence of mitochondrial membrane potential formation in the presence of an H₂S scavenger suggests that free H₂S is not the primary bioenergetic substrate (Fig. S4G, H). Instead, these data support the notion that hydropersulfides and polysulfides, likely generated downstream of H₂S, represent the functionally relevant substrates for SQR. In this context, hydropersulfides and polysulfides appear to be more efficient electron donors to the ETC than H₂S itself, thereby more effectively supporting mitochondrial membrane potential formation and bioenergetics. This interpretation is consistent with our metabolomic data showing accumulation of persulfide species in SQR-deficient systems (Fig. 5 and S2) and highlights a central role of supersulfide species in driving sulfur respiration.

In this context, the recent development of chemical supersulfide boosters provides an important proof-of-concept for the therapeutic targeting of sulfur respiration. For example, 2*H*-thiopyran-2-thione sulfine has been shown to convert H_2_S into hydrogen persulfide and elevate intracellular supersulfide levels, leading to enhanced mitochondrial membrane potential and energy metabolism [37]. These findings suggest that the pharmacological activation of SQR-mimic sulfur metabolic pathways may represent a promising strategy to augment mitochondrial bioenergetics, particularly in conditions of mitochondrial dysfunction.

Of note, sulfur respiration does not require a functional complex I, and electrons and protons enter from CoQ and proceed to the ETC. This suggests that the dietary supplementation of cystine activates sulfur respiration and increases ATP production in patients with mitochondrial diseases, particularly those suffering from complex I deficiency. A recent paper stated that the Leigh syndrome mouse model, which possesses complex I deficiency due to the Ndufs4 deletion, is rescued by brain hypoxia [38]. On the basis of our current study, we hypothesize that hypoxia may drive a compensatory induction of sulfur respiration that enables energy production independent of complex I. If this is the case, activation of mitochondrial sulfur metabolism may be a potential therapeutic strategy for mitochondrial diseases with complex I defects as it may stimulate mitochondrial energy metabolism. Promotion of sulfur respiration, besides aiding in the treatment of mitochondrial diseases, would benefit the maintenance of healthy tissues and organs with a relatively high demand for energy, such as the skeletal muscle, liver, and possibly neural tissues. We thus envision the exciting possibility of an era of novel preventative and therapeutic approaches in various diseases as well as in the control of aging and longevity.

In recent years, several reports have described the beneficial effects of hydrogen sulfide on enhancing longevity [39–41]. Our current findings provide mechanistic insights into these observations by demonstrating that the lifespan-extending effects of hydrogen sulfide and hydropersulfides are strictly dependent on the presence of functional SQR. In the absence of SQR, neither NaHS nor Na₂S₂ treatment was sufficient to extend the yeast lifespan, highlighting the essential role of SQR in converting sulfide species into bioenergetically useful reducing equivalents. Our data suggest that sulfide-mediated lifespan extension involves mitochondrial bioenergetics and is not a direct antioxidant effect of sulfide or its derivatives. Moreover, our results imply that the beneficial effects of exogenous sulfur-based donors are not merely due to a general redox modulation but rather involve specific enzymatic oxidation processes mediated by SQR, which in turn support mitochondrial membrane potential and ATP production. This mechanism likely underlies the evolutionarily conserved role of hydrogen sulfide in promoting longevity across different species.

In conclusion, our findings strengthen the emerging view that mitochondrial sulfur metabolism, particularly the SQR-dependent oxidation of persulfide species, constitutes a fundamental and physiologically relevant form of energy production in eukaryotic cells.

## 4. Materials and Methods

### 4.1. Yeast strain, medium, and plasmid

The media used for the growth were Yeast Extract-Peptone-Dextrose (YPD), Synthetic Complete (SC), and Sehgal and Gibbon’s (SG). YPD contains 1% yeast extract (Difco Laboratories, Detroit, MI), 2% peptone (Difco Laboratories), and 2% glucose. SC has 2% glucose, 0.67% yeast nitrogen base without amino acids (Difco Laboratories), and 0.2% dropout supplement (Takara Bio, Shiga, Japan). SG contains 2% galactose instead of glucose in SC. The appropriate amino acids and/or nucleotides were removed for auxotrophic markers. When necessary, 2% agar (Nacalai Tesque, Kyoto, Japan) was added to solidify the medium.

The yeast *Schizosaccharomyces pombe* with a 972 background was used in this study. The *HMT2* disrupted strain (*Δhmt2*) was constructed by using a two-step PCR method with minor modifications [42]. The 5,000-bp homologous sequences were added to the kanMX6 deletion cassette by using a two-step overlapping PCR with primers and pFA6-KanMX6 plasmid. The PCR fragment was integrated into the genome by transformation via the LiAc/SS carrier DNA/PEG method [43]. Screening for G418 resistance was performed, and the correct integration event was verified via PCR with chromosomal DNA.

### 4.2. Growth inhibition by CdCl_2_ and NaHS in yeast

Yeast cells were cultured in synthetic complete medium [SC, 2% glucose, 0.67% yeast nitrogen base without amino acids, and 0.2% dropout supplement (Takara Bio Inc., Shiga, Japan)] at a starting OD_600_ of 0.01. After 10 min at 30 °C, CdCl_2_ (10-300 μM) or NaHS (50-1,000 μM) was added to the culture, and the culture was allowed to continue growing at 30°C. The OD_600_ of the culture was measured after 24 h. The growth rate was converted into percent growth relative to the untreated controls.

### 4.3. Growth test of yeast on glucose and glycerol media

For the agar plate culture, yeast cells were grown at 30 °C in YE (3% glucose, 0.5% yeast extract) to the stationary growth phase and then spotted in 10-fold dilutions at a starting OD_600_ of 1.0 on YE or YEGly medium (3% glycerol, 0.1% glucose, 0.5% yeast extract) containing 2% agar. The plates were then incubated at 30 °C for 2 days. For the liquid culture, yeast cells were cultured at 30 °C in YE or YEGly medium at a starting 0.1 of OD_600_. Cell growth was monitored by measuring OD_600_.

### 4.4. Measurement of mitochondrial membrane potential of yeast

The mitochondrial membrane potential was measured by using JC-1 (Molecular Probes, Eugene, OR, USA), a cationic dye that shows potential-dependent accumulation in the mitochondria [44]. Yeast cells were cultured in SC medium at 30 °C for 24 h (log phase) or 48 h (stationary phase), harvested by centrifugation, and washed once with phosphate-buffered saline (PBS) buffer. The harvested cells were incubated with 10 μM JC-1 in PBS for 30 min at 30°C and then subjected to flow cytometry analysis. Flow cytometry was performed by using a BD Accuri C6 flow cytometer (BD Bioscience, Franklin Lakes, NJ, USA), and the FL2/FL1 values of individual cells were calculated to determine mitochondrial membrane potential. Negative control cells with depolarized mitochondrial membranes were treated with 5 μM carbonyl cyanide m-chlorophenyl hydrazine (CCCP) (Sigma-Aldrich, St. Louis, MO) for 30 min at room temperature before JC-1 labeling.

### 4.5. Determination of intracellular ATP levels of yeast

Cultured cells of about 1 × 10^7^ cells (OD = 1.0) were harvested, washed three times with ice-cold PBS, and resuspended in 200 μL of sterilized water. After being frozen by liquid nitrogen, the freezing cells were heated for 10 min at 95 °C to release ATP to the outside of the cells. ATP levels in 100 μL of the supernatant were determined by IntraCellular ATP assay kit (Toyo B-net, Tokyo, Japan).

### 4.6. Supersulfide metabolome analysis of yeast

Supersulfides and other sulfur metabolites produced by SQR-deficient fission yeast were analyzed by using liquid chromatography-electrospray ionization-tandem mass spectrometry (LC-ESI-MS/MS) according to our previously reported method [6, 7]. For analyses of intracellular supersulfide and other sulfur metabolites levels, the cultured yeasts were lysed by sonication in a cold methanol solution containing 1 mM β-(4-hydroxyphenyl)ethyl iodoacetamide (HPE-IAM), after which cell lysates were incubated for 20 min at 37 °C. After centrifugation, aliquots of the supernatants of the cell lysates were diluted 20 times with 0.1% formic acid containing known amounts of isotope-labeled internal standards, which were then analyzed via LC-ESI-MS/MS (LCMS-8060NX; Shimadzu) for supersulfide determination (Fig. 2).

### 4.7. Chronological lifespan assay

Chronological lifespan was determined according to a previously described method with simple modifications [36, 45]. Briefly, yeast cells were cultured in SC medium (at a starting OD_600_ of 0.1) at 30 °C with shaking. Measurement of cell viability began after 72 h of culture (day 0) and continued every 2-3 days by plating a fraction of the culture onto a fresh YPD plate to determine the number of colony-forming units (CFUs). To investigate the effects of NaHS and Na_2_S_2_, the donor compounds were added every 3 days. Cell survival rates were calculated by normalizing the CFUs at each time point to the CFUs obtained at day 0.

### 4.8. Establishment of immortalized MEFs

*Sqrdl*^ΔN/ΔN^ mice were generated as previously reported [14]. WT and *Sqrdl*^ΔN/ΔN^ iMEFs were established from mouse embryos at E13.5 and immortalized by lentiviral introduction of SV40 large T antigen. WT and *Sqrdl*^ΔN/ΔN^ embryos were obtained from *Sqrdl*^ΔN/+^ pregnant females mated with *Sqrdl*^ΔN/+^ males. Stable transformants were selected via 2 μg/ml puromycin, and three independent WT iMEF lines and *Sqrdl*^ΔN/ΔN^ iMEF lines were established.

### 4.9. Measurement of mitochondrial membrane potential of iMEFs

To determine the membrane potential of mitochondria under several experimental conditions, JC-1 staining was performed according to the method previously reported [43, 46]. Accumulation of the cell-permeable JC-1 probe (Abcam) in the mitochondria depends on the membrane potential and is associated with a fluorescence emission shift from green (emission 527 nm) to red (emission 590 nm). Briefly, WT and *Sqrdl*^ΔN/ΔN^ iMEFs, cultured in 8-well multi chamber slides (Matsunami, Osaka, Japan) coated with collagen, were treated with mitochondrial ETC inhibitors (rotenone, atpenin A5, and antimycin A) and ADP/ATP carrier inhibitor CATR [33] as described in the figure legends. For JC-1 staining, cultured cells were washed with HKRB buffer (20 mM HEPES, 103 mM NaCl, 4.77 mM KCl, 0.5 mM CaCl_2_, 1.2 mM MgCl_2_, 1.2 mM KH_2_PO_4_, 25 mM NaHCO_3_, and 15 mM glucose, pH 7.3), incubated with 20 µM JC-1 at 37 °C for 20 min, rinsed twice with HKRB buffer, and examined with a confocal laser scanning microscope (Nikon C2 plus, with NIS-Elements 5.01 software). ImageJ software was used for image processing and calculation of JC-1 fluorescence intensities.

### 4.10. Measurement of the OCR of WT and mitochondria-specific SQR-deficient iMEFs

The OCR of mitochondria from WT and mitochondria-specific *Sqrdl*-deficient iMEFs was measured by using the XF96 Extracellular Flux Analyzer (Agilent). The OCR was determined under various conditions (oligomycin, dinitrophenol, and rotenone plus antimycin A). Using these metabolic modulators allows multiple parameters of mitochondrial function to be determined, as previously reported [47]. Oligomycin was used to inhibit ATP synthase, 2,4-dinitrophenol (DNP) to uncouple mitochondria and yield maximal OCR, and rotenone plus antimycin A to inhibit mitochondrial oxygen consumption.

### 4.11. Measurement of the membrane potential of isolated mitochondria

iMEFs mitochondria were isolated by differential centrifugation. WT and *Sqrdl*^ΔN/ΔN^ iMEFs were homogenized in the extraction buffer (10 mM Tris-HCl, 250 mM sucrose, and 0.5 mM EGTA, pH 7.4) with a Dounce homogenizer (Teflon glass) and were centrifuged at 2,000 x g at 4 °C for 5 min to remove nuclei and cell debris. The supernatants were centrifuged at 12,000 x g at 4 °C for 10 min, and mitochondrial pellets were washed three times with a buffer and kept at -80 °C until use. To determine the membrane potential in isolated mitochondria under several experimental conditions, TMRE staining was performed according to the method previously reported [48]. The TMRE fluorescence intensity at a maximum emission of 574 nm depends on the mitochondrial membrane potential. Briefly, mitochondria from WT and *Sqrdl*^ΔN/ΔN^ iMEFs were adsorbed on the bottom of 8-well slides (Ibidi, Martinsreid, Germany) coated with poly-D-lysine (Sigma-Aldrich, St. Luis, MO, USA), after which they were stained with 10 nM TMRE (Biotium, Fremont, CA, USA) and 50 nM MitoTracker Green FM (Thermo Fisher Scientific, Waltham, MA, USA) at 25 °C for 10 min. TMRE fluorescence intensity (561 nm excitation and 605 ± 15 nm emission), which is positive with MitoTracker Green FM fluorescence, was measured in individual mitochondria by using the region of interest (ROI) tool of the confocal laser scanning microscope (Nikon C2 plus, with NIS elements version 5.01 software). ROIs, 200 each for WT and *Sqrdl*^ΔN/ΔN^ mitochondria, were manually drawn around a single mitochondrion that did not overlap with others, and the average non-zero pixel intensity within the ROIs was measured and plotted against time as arbitrary units. Because damage to some mitochondria is technically inevitable, we selected mitochondria that showed a sufficient response to TMPD/ascorbate. Specifically, of 200 ROIs, the top 30 ROIs were selected for analysis of TMRE fluorescence intensity in response to TMPD/ascorbate, and their means and s.d. values were calculated.

### 4.12 Measurement of the OCR in liver mitochondria

#### 4.12.1. Isolation of liver mitochondria

Mitochondria were isolated from 4- to 6-week-old mice by using two-step differential centrifugations, as previously described [49] with minor modifications. In brief, after gallbladder removal, fresh liver specimens weighing 1.0-1.5 g were minced and homogenized in ice-cold isolation buffer (10 mM Tris/MOPS, 1 mM EGTA/Tris, and 200 mM sucrose, pH 7.4). When liver weight was less than 1 g, livers were pooled. Homogenates were centrifuged at 600 x *g* and 4 °C for 10 min to remove cellular debris and nuclei. Supernatants containing mitochondrial fractions were then centrifuged at 7,000 x g at 4 °C for 10 min. Obtained mitochondrial pellets were washed with a new isolation buffer and underwent centrifugation again (7,000 x g for 10 min). Supernatants were discarded, and the final mitochondrial pellet was resuspended in a remaining minimal volume of the isolation buffer. The protein concentration was determined with the BCA Protein Assay Kit (Thermo Fisher Scientific). Absorbance at 562 nm was measured with NanoDrop 2000 (Thermo Fisher Scientific).

#### 4.12.2. Measurement of liver mitochondrial respiratory function

Mitochondrial oxygen consumption was measured by using a Clark-type electrode (Oxytherm system; Hansatech Instruments, Norfolk, UK). All experiments were conducted with isolated mitochondria, 1 mg protein/ml in experimental buffer (125 mM KCl, 10 mM Tris/MOPS, 10 mM EGTA/Tris, and 1 mM Pi, pH 7.4), with continuous stirring at 25 °C. The OCR through complex I was measured as a mitochondrial basic function by sequentially adding 2.5 mM malate/5 mM glutamate, 150 µM ADP, and 100 nM carbonyl cyanide 4-(trifluoromethoxy) phenylhydrazone (FCCP). The respiratory control index (RCI) was calculated by using the ratio of state 3 to state 4. The electron donation ability of SQR was analyzed by three additions of 10 µM NaHS in the presence of 100 nM FCCP, which was followed by addition of 250 µM TMPD/2.5 mM ascorbic acid for evaluation of complex IV integrity. Then, 100 µM sodium cyanide (NaCN) was added to terminate the electron transfer.

### 4.13. Quantification of supersulfides

Yeast, and iMEFs were analyzed for the low molecular weight analytes cysteine (CysSH), cysteine hydropersulfide (CysSSH), glutathione (GSH), GSSH, bis-S (H_2_S), bis-SS (HSSH), cystine (CysSSCys), oxidized glutathione or glutathione disulfide (GSSG), GSSSG, and thiosulfate (HS_2_O_3_^-^) as previously described [6]. Briefly, samples were homogenized with a Psychotron homogenizer (Microtec Co. Ltd.) or sonicator in ice-cold methanol in the presence of 5 mM HPE-IAM. Homogenates were incubated at 37 °C for 20 min and then centrifuged at 15,000 rpm at 4 °C for 10 min. Supernatants were separated and diluted 10 times with 0.1% formic acid. Stable isotope-labeled internal standards dissolved in 0.1% formic acid solution were added to the samples to a final concentration of 25 nM. A sample of 10 µl was injected into the LC-ESI-MS/MS instrument (LCMS-8060). Various persulfide derivatives were identified and quantified by means of MRM, as previously described [6]. Cell pellets were reconstituted in 0.1% SDS in PBS buffer, homogenized by sonication, and centrifuged again at 15,000 rpm at 4 °C for 10 min. The BCA protein assay was performed with the supernatants to determine the total protein content of the samples. Each analyte concentration was normalized to the measured protein concentration.

Isolated mitochondria (0.45 mg/ml) from iMEFs were treated with 50 µM NaHS and 100 µM GSSSG, or were not treated, at 37 °C for 60 min and 15 min, respectively. After incubation, supersulfides were extracted with 5 mM HPE-IAM containing methanol and subjected to LC-ESI-MS/MS analysis by following the procedure described above.

### 4.14. Statistical analysis

Statistical significance was evaluated by using an unpaired two-sample Student’s *t*-test for two-group comparisons and one-way/two-way ANOVA with Tukey’s test for multiple-group comparisons. Kaplan-Meier analysis was used for survival of *Sqrdl*^ΔN/ΔN^ mice, and statistical significance was evaluated via the log-rank test. These analyses were performed by using Microsoft Office Excel (Microsoft) and Prism 9 (GraphPad Software, Inc., San Diego, CA, USA). *P* < 0.05 was considered to be statistically significant.

## Acknowledgments

This work was supported in part by Transformative Research Areas, International Leading Research, Scientific Research [(S), (B), (C), Challenging Exploratory Research] from the Ministry of Education, Culture, Sports, Science and Technology (MEXT), Japan, to T. A. (21H05263, 22K19397 23K20040 and 24H00063), T. M. (22K06893, 25K03490), S. O. (23K14333), T. T. (23K06094) and M. M. (23K06145); by Japan Science and Technology Agency, Japan, CREST Grant Number JPMJCR2024, to T. A.; and by a grant from the Japan Agency for Medical Research and Development (AMED) to T. A. (JP21zf0127001). M.S. is supported by a NIH F31 Predoctoral Fellowship (F31HL170516). J.Y. is supported by the WISE Program for Sustainability in the Dynamic Earth of Tohoku University.

## Author Contributions

J.Y., T.M., A.N., M.S., T.I., M.J., S.O., T.T., U.B., H.M., M.M., and T.A. designed research; J.Y., T.M., A.N., M.S., T.I., M.J., S.O., T.T., U.B., and M.M. performed experiments; J.Y., T.M., A.N., M.S., M.M., and T.A. wrote the paper.

## Supplementary Information

**Supplementary Fig. 1.**
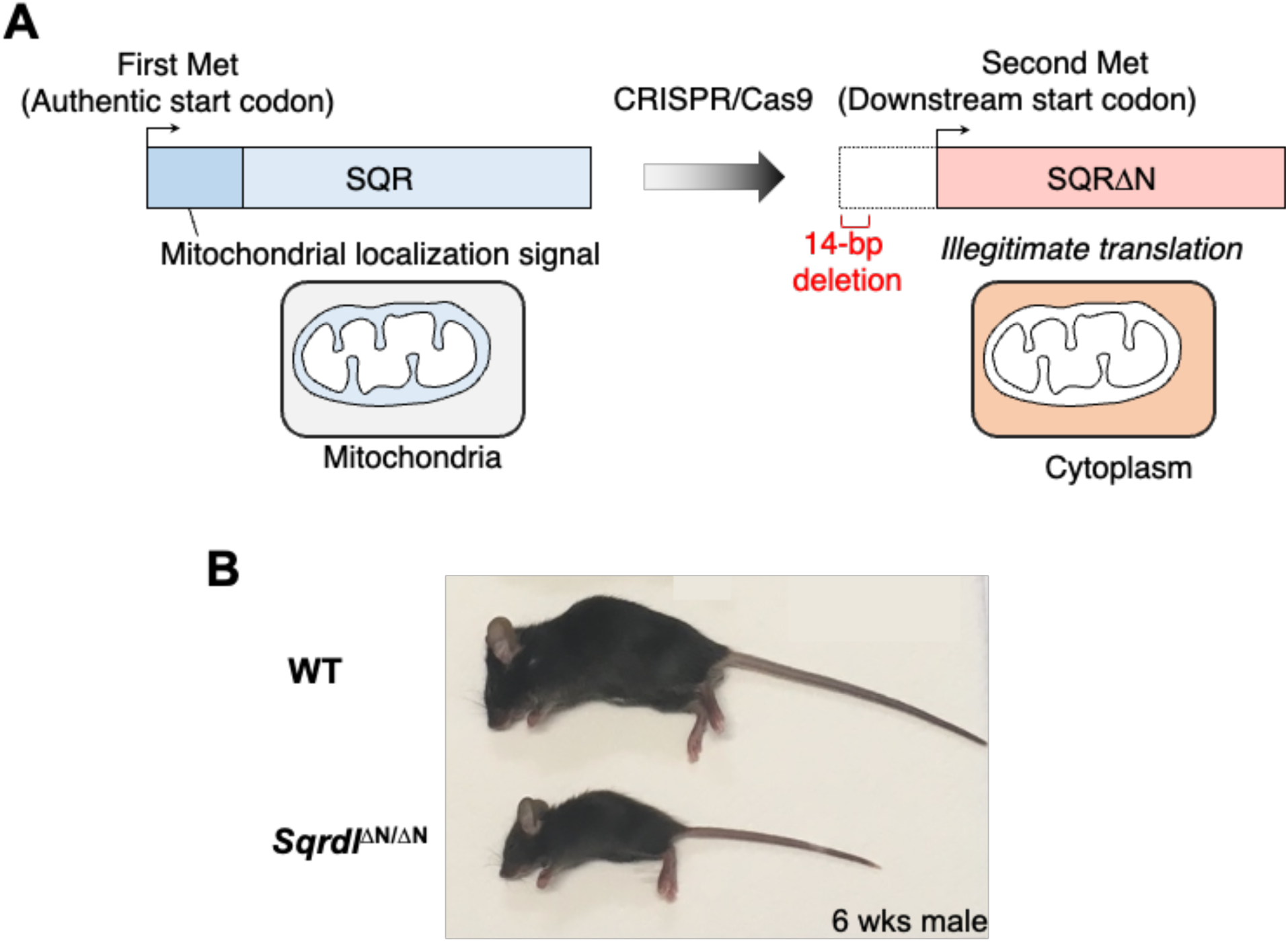
Generation of SQR mutant mice. (A) Schematic presentation of SQR proteins expressed by WT and SQRΔN alleles. (B) Macroscopic appearance of 6-week-old WT, and *Sqrdl*^ΔN/ΔN^ littermate male mice.

**Supplementary Fig. 2.**
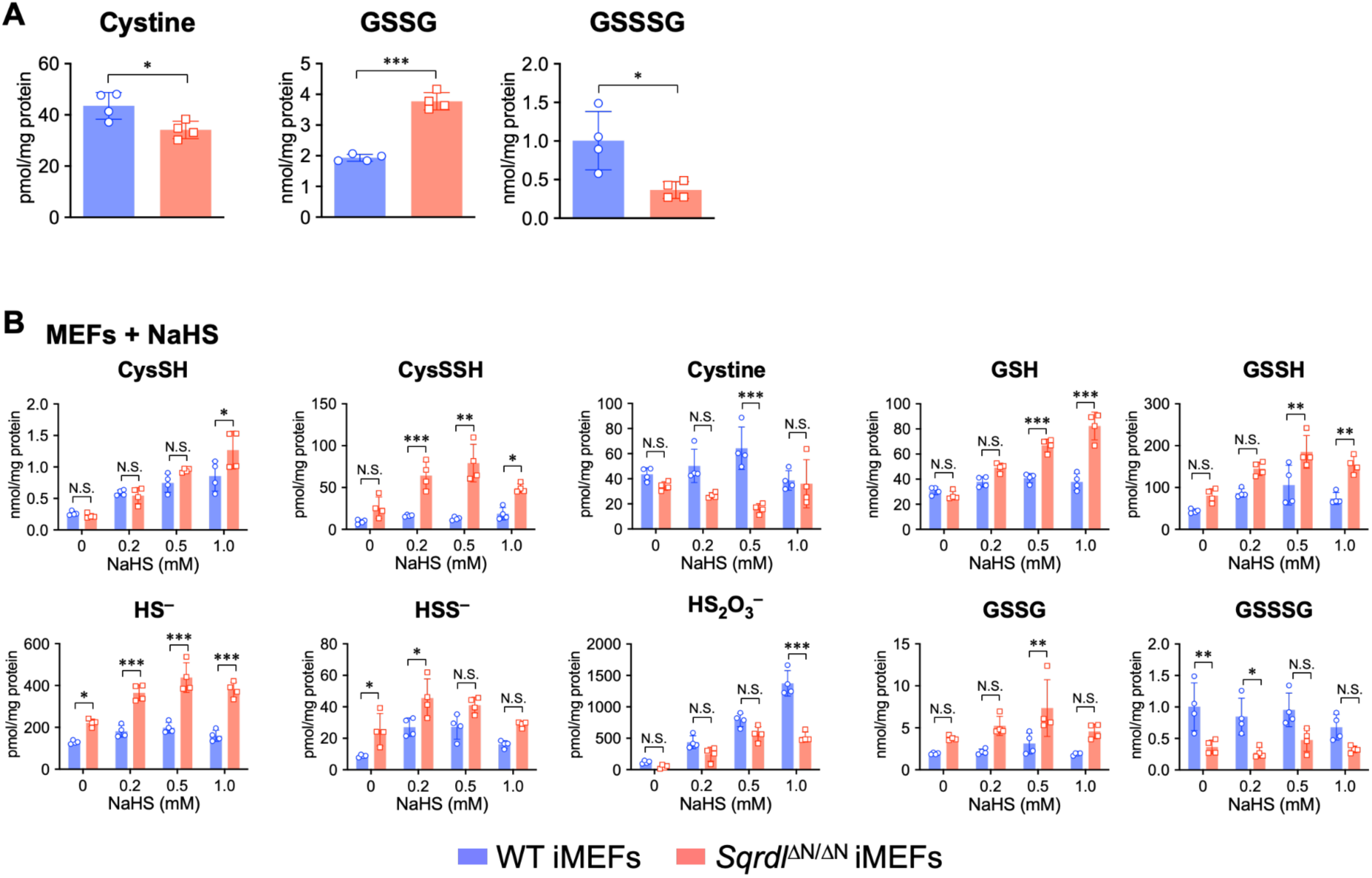
Sulfur metabolome analysis of WT and *Sqrdl*^ΔN/ΔN^ iMEFs. (A,B) Quantification of sulfur metabolites in (A) iMEFs, (B) iMEFs treated with NaHS or without such treatment. Data are represented as mean ± s.d. (*n* = 3-5). \**P* < 0.05, \*\**P* < 0.01, and \*\*\**P* < 0.001, determined by Student’s *t*-test.

**Supplementary Fig. 3.**
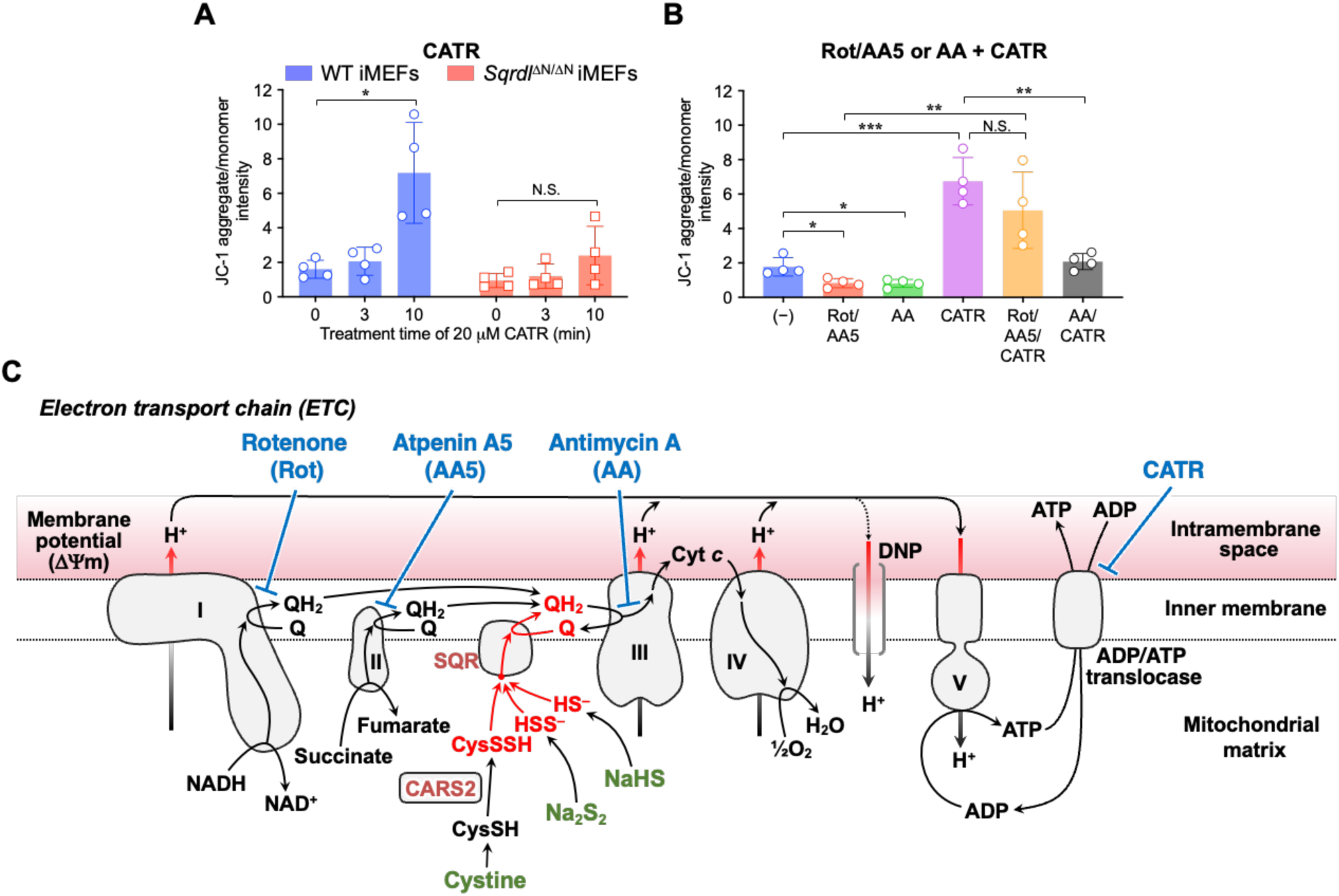
(A) Response of mitochondrial membrane potential to CATR in WT and *Sqrdl*^ΔN/ΔN^ iMEFs. (B) Effects of the addition of rotenone (Rot), atpenin A5 (AA5), antimycin A (AA), and CATR on mitochondrial membrane potential of WT iMEFs. Data are mean ± s.d. (n = 3 or 4). \**P* < 0.05, \*\**P* < 0.01, \*\*\**P* < 0.001, N.S. (not significant), determined by one-way ANOVA with Tukey’s test. (C) Schematic drawing of the ETC in mitochondria with SQR and CARS2. Inhibitors and substrates used in experiments indicated in Fig. 7 appear in blue and green, respectively. Cyt *c*, cytochrome c; ΔΨm, membrane potential.

**Supplementary Fig. 4.**
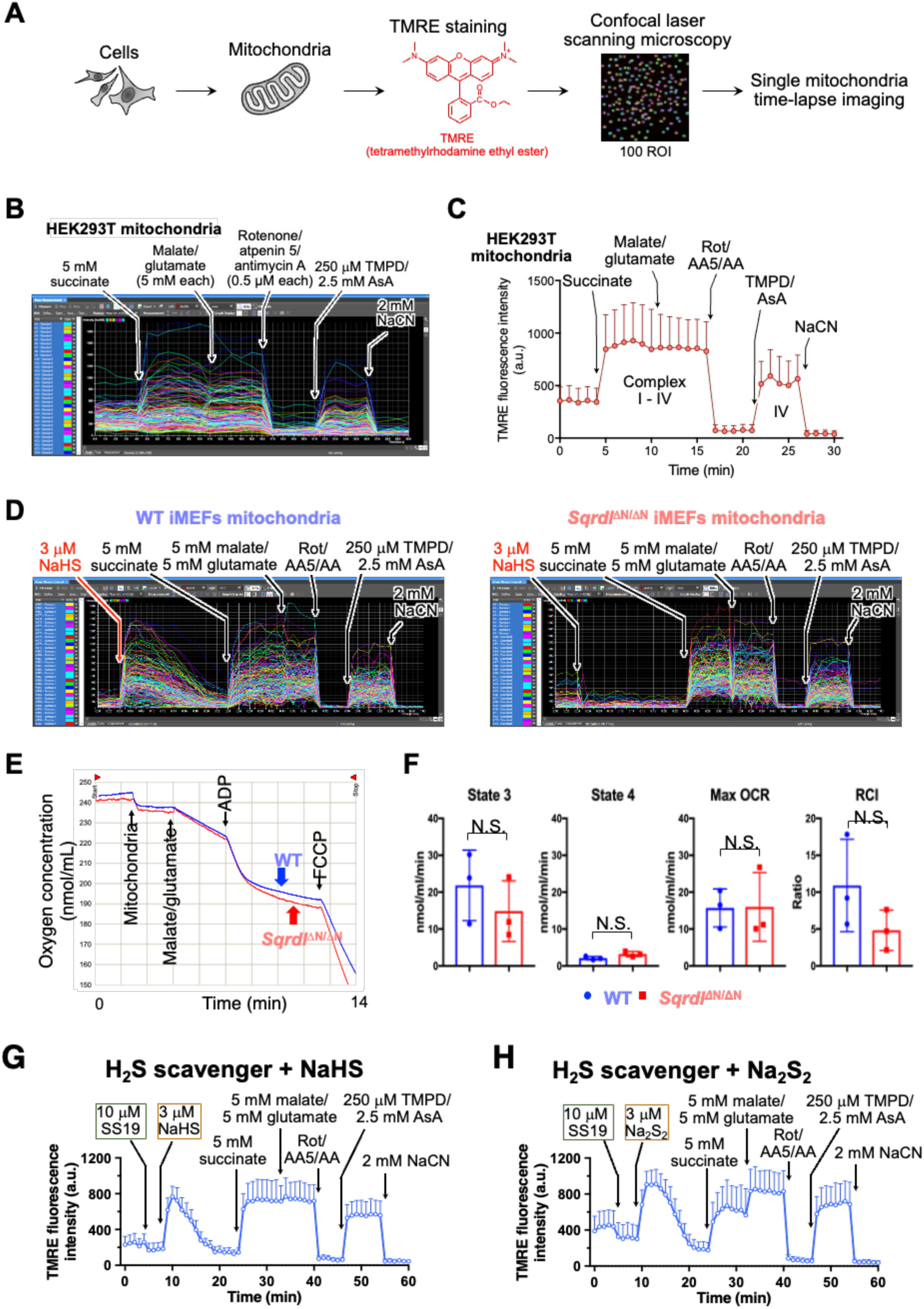
Evaluation of the SQR contribution to mitochondrial activity in isolated mitochondria. (A-C) Scheme and representative images of time-lapse changes in TMRE fluorescence of mitochondria isolated from HEK293T cells. (D) Representative images of time-lapse changes in TMRE fluorescence of mitochondria isolated from WT and *Sqrdl*^ΔN/ΔN^ iMEFs. (E) Representative oxygen consumption curves of mitochondria isolated from the livers of WT and *Sqrdl*^ΔN/ΔN^ mice. (F) OCR and RCI of WT and *Sqrdl*^ΔN/ΔN^ mitochondria. OCR at states 3 and 4 and the maximum OCR with uncoupler FCCP from three independent experiments. The RCI was obtained from the ratio of state 3 OCR to state 4 OCR. (G, H) NaHS-induced (G) and Na_2_S_2_-induced (H) membrane potential in mitochondria isolated from WT iMEFs in the presence of 10 µM SS19, a hydrogen sulfide scavenger. Data are represented as mean ± s.d.

